# A hyperpolarizing neuron recruits undocked innexin hemichannels to transmit neural information in *Caenorhabditis elegans*

**DOI:** 10.1101/2023.08.21.554224

**Authors:** Airi Nakayama, Masakatsu Watanabe, Riku Yamashiro, Hiroo Kuroyanagi, Atsunori Oshima, Ikue Mori, Shunji Nakano

## Abstract

While depolarization of neuronal membrane is known to evoke the neurotransmitter release from synaptic vesicles, hyperpolarization is regarded as a resting state of chemical neurotransmission. Here we report that hyperpolarizing neurons can actively signal neural information by employing undocked hemichannels. We show that UNC-7, a member of the innexin family in *Caenorhabditis elegans,* functions as a hemichannel in thermosensory neurons and transmits temperature information from the thermosensory neurons to their post-synaptic interneurons. By monitoring neural activities in freely behaving animals, we find that hyperpolarizing thermosensory neurons inhibit the activity of the interneurons and that UNC-7 hemichannels regulate this process. UNC-7 is required to control thermotaxis behavior and functions independently of synaptic vesicle exocytosis. Our findings suggest that innexin hemichannels mediate neurotransmission from hyperpolarizing neurons in a manner that is distinct from the synaptic transmission, expanding the way of neural circuitry operations.

## Introduction

Neurotransmission is a primary means of neural communication and plays an essential role in the generation of animal behaviors. A common form of neural communication is chemical synaptic transmission^1,2^, in which the neurotransmitters are released through synaptic vesicle exocytosis. This exocytosis occurs in response to calcium influx at the presynaptic site, which is triggered by membrane depolarization that causes voltage-gated calcium channels to open. Most chemical transmission is induced by action potentials, and these action potential-dependent transmissions do not occur in hyperpolarizing neurons. Likewise, at neurons with graded membrane potential, the synaptic transmitter secretes tonically, such that neurotransmitter release is increased by depolarization and decreased by hyperpolarization^3^.

Neural communication is also mediated by electrical synapses, also known as gap junctions^4,5^. The core proteins of gap junctions, connexins in vertebrates and innexins in invertebrates^6,7^, form undocked membrane channels, hemichannels, which on their own can promote the passage of small molecules between the cytosol and extracellular space^8,9^. Gap junction channels are assembled by docking of two opposing hemichannels from adjacent cells, thereby connecting the cytoplasms of adjacent neurons and forming electrical coupling that can mediate neuronal synchrony^4,5^. For instance, gap junctions synchronize neuronal activities in inferior olivary nucleus through propagating depolarizing spikes^10,11^.

In addition to depolarization, electrical synapses are also reported to spread hyperpolarizing potentials. In the Golgi neurons of mammalian cerebellum, gap junctions composed of connexins were shown to facilitate the propagation of hyperpolarization after a brief depolarizing current^12,13^. This observation implies that hyperpolarizing neurons might also be engaged in controlling the dynamics of neural circuitry. However, the mechanisms underlying neurotransmission from hyperpolarizing neurons and their roles in animal behaviors have remained largely unexplored.

The *C. elegans* thermotaxis behavior provides an effective model in which to address the mechanisms of neurotransmission from hyperpolarizing neurons. The temperature preference of *C. elegans* is plastic and is determined by the past experience: the animals prefer the temperature at which they have been previously cultivated in the presence of food^14^. When placed on a temperature gradient, the animals migrate toward the cultivation temperature^15–17^. In light of the complete knowledge of the *C. elegans* connectome composed of only 302 neurons^18,19^, the neural circuitry regulating thermotaxis have been extensively studied^15,20,21^. Central to this circuit is the AFD thermosensory neuron essential for temperature sensing^15^. In response to warming stimuli, the AFD thermosensory neuron depolarizes and increases its intracellular calcium concentration^16,22–26^. The depolarized AFD neurons control the activity of their post-synaptic partner, the AIY interneuron^20,27,28^. Regulation of the AIY activity by AFD is bidirectional, and the AIY response represents stimulus valence^29,30^: AIY is excited in response to warming stimulus with the positive valence where the temperature is increased toward the cultivation temperature, while the AIY activity is inhibited upon warming above the cultivation temperature^30^.

By contrast, very little is known about how the AFD neurons transmit neuronal outputs in response to cooling stimuli. Previous studies demonstrated that cooling stimuli hyperpolarized the AFD membrane potentials^24^ and that the AFD neuron is indispensable for the behavioral control upon temperature cooling^16,17^. However, how the hyperpolarized AFD neurons transmit neural information and control the neural circuitry to generate appropriate behaviors remains elusive.

In this study, we identify *unc-7*, which encodes an innexin protein, as a new regulator for thermotaxis and show that UNC-7 acts as a hemichannel in AFD to transmit temperature information to the AIY interneurons. The UNC-7 hemichannels regulate AIY neuronal activity upon membrane hyperpolarization of the AFD neuron. Intriguingly, UNC-7 controls thermotaxis in parallel to the chemical transmission from AFD. Our findings suggest that hyperpolarized neurons can actively transmit neural information by employing innexin hemichannels that function independently of synaptic vesicle exocytosis.

## Results

### UNC-7 acts in the AFD thermosensory neurons to regulate the *C. elegans* thermotaxis

We have previously shown that *inx-4*, a member of innexins, functions in the AFD thermosensory neurons to regulate thermotaxis^31^. In this study, we conducted a genome-wide survey of innexins, aiming to reveal the roles of the innexin family genes in the control of thermotaxis (Fig. 1). Of the 24 remaining innexin genes present in the *C. elegans* genome, we focused on the genes that are dispensable for viability and locomotion and are expressed in the nervous system^32–35^. We examined the thermotaxis behaviors of mutants for such innexin genes and found that all innexin mutants examined displayed the wild-type thermotaxis phenotype (Fig. 1b). We also assessed the effect of AFD-specific overexpression of the innexin genes. Seven innexin genes - *inx-1, inx-2, inx-4, inx-7, inx-10, inx-19* and *unc-7* - were reported to be expressed in AFD of the wild-type animals^36^. We overexpressed each of these genes in AFD and observed that overexpression of *unc-7* caused a thermophilic phenotype (Fig. 1c). While the wild-type animals cultivated at 20 °C stayed around the cultivation temperature, animals overexpressing UNC-7 in AFD migrated toward a temperature region higher than the cultivation temperature (Fig. 1c6). By contrast, animals overexpressing other innexin genes in AFD did not show abnormality in thermotaxis.

**Figure 1.**
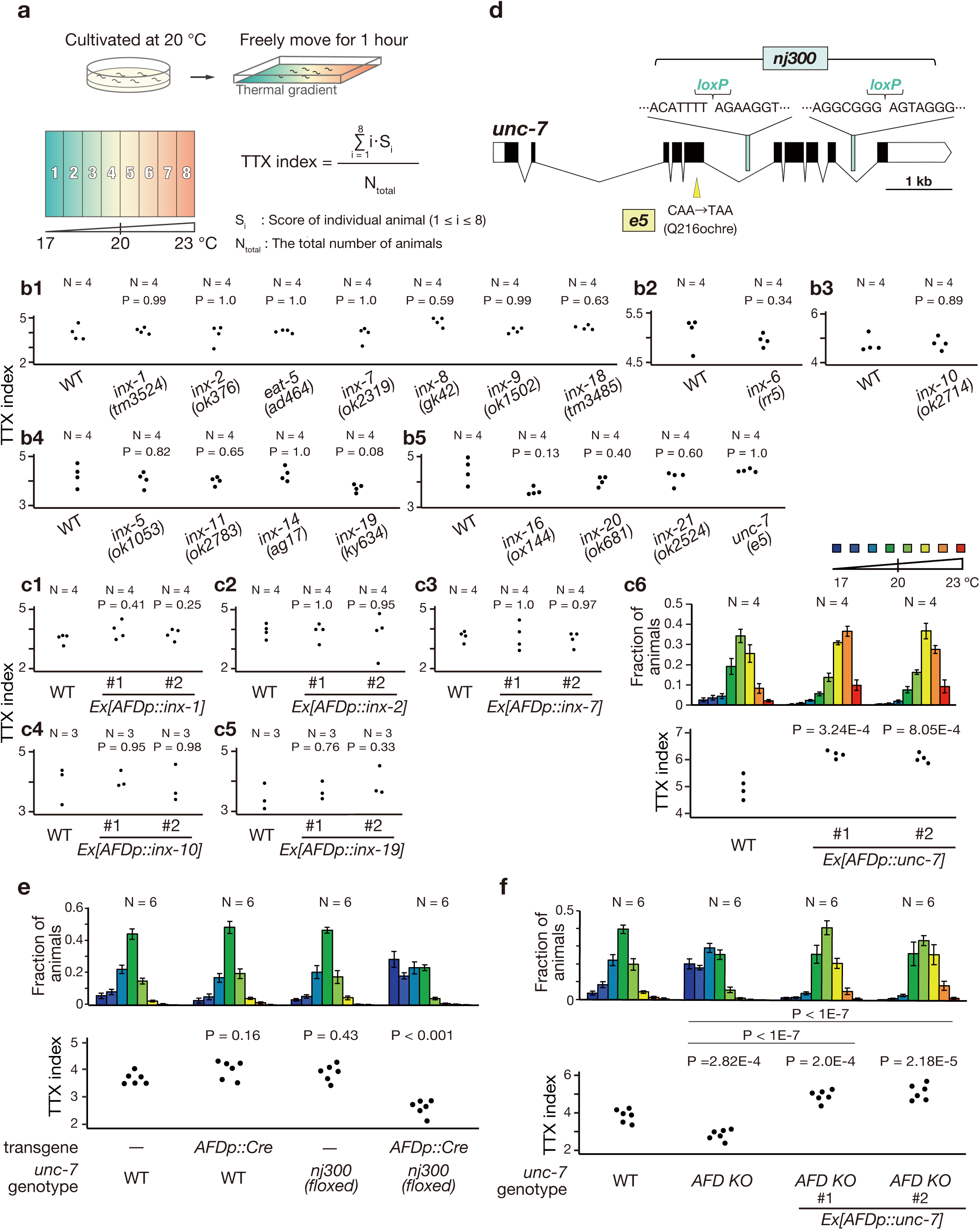
UNC-7 functions in the AFD thermosensory neuron to regulate the thermotaxis behavior. (a) Schematic of thermotaxis (TTX) assay and the formulas for TTX index to quantify the thermotaxis behavior (Materials and Methods). (b) TTX behaviors of innexin single mutants. TTX indexes are shown in the dot plots. P-values were determined by Dunnet test (b1 and b4), by Exact Wilcoxon rank sum test (b2 and b3) or by Steel test (b5). (c) TTX behaviors of animals overexpressing innexin genes in the AFD thermosensory neuron. Distributions of the animals on the thermal gradients of the TTX plate are displayed in the histograms. The fraction of the population in each temperature section is shown as mean ± SEM. TTX indexes are shown as in (b). P-values were determined by Steel test (c1) or by Dunnet test (c2-c6). (d) Gene structure of *unc-7 (isoform a)* is shown. Black boxes, white boxes and lines indicate exons, untranslated regions and introns respectively. Yellow triangle indicates the mutation site of *unc-7(e5)*. Blue squares indicate the genome sequences flanking the *loxP* insertion sites in *unc-7(nj300, floxed)*. (e) TTX behavior of *unc-7(AFD KO)* animals lacking *unc-7* specifically in the AFD neuron. P-values were determined by Dunnet test. (f) TTX behavior of *unc-7(AFD KO)* animals carrying a transgene that expresses the wild-type *unc-7* gene specifically in the AFD neuron. P-values were determined by 1-way ANOVA with Tukey HSD test.

Mutants harboring a presumptive null allele of *unc-7, unc-7(e5)*, were uncoordinated^32^, and hence we could not test their thermotaxis behavior. To address whether a loss of *unc-7* function affects thermotaxis, we attempted to generate strains lacking *unc-7* only in AFD using the Cre/*loxP* system. We inserted two *loxP* sequences into the *unc-7* locus and crossed with this *unc-7(nj300)* animal a strain carrying a single-copy insertion of the *Cre* gene fused with the AFD-specific *gcy-8L* promoter (Fig. 1d). This strain, hereafter referred to as *unc-7(AFD KO)* animals, displayed a cryophilic defect and migrated toward a lower temperature region than the wild-type animals did (Fig. 1e). Strains carrying *unc-7(nj300)* or the *gcy-8Lp::Cre* insertion alone did not show abnormality in thermotaxis. The thermotaxis defect of *unc-7(AFD KO)* was rescued by expressing *unc-7* specifically in AFD (Fig. 1f). These results indicate that UNC-7 functions in AFD to regulate thermotaxis.

### UNC-7 is required later than the larval stage to regulate thermotaxis

A previous study showed that UNC-7 is required for presynaptic differentiation during synaptogenesis^37^. This observation suggests that the thermotaxis defect of *unc-7(AFD KO)* could be attributable to abnormalities of synaptogenesis of AFD. To assess whether UNC-7 is required for the development of the AFD thermosensory neurons, we examined the critical period of UNC-7 for the regulation of thermotaxis by Auxin Induced Degradation (AID) system^38^. We expressed TIR1 in AFD of animals in which *unc-7* was tagged with a degron sequence. While the animals cultivated with auxin throughout the development showed cryophilic defects, the animals cultivated without auxin did not show abnormality in thermotaxis (Supplementary Fig. 1a, b), indicating that the AID system successfully knocked down the UNC-7 activity. When the animals were cultivated in the absence of auxin for two days, at which most animals had grown to the second or third larval stage and were then transferred onto plates with auxin, they displayed a cryophilic defect (Supplementary Fig. 1c, d). Conversely, the animals treated with auxin only for the first two days of their development did not exhibit thermotaxis defects. These results indicated that the activity of UNC-7 in AFD is required at or later than the third larval stage, the period at which AFD has connected to the majority of its postsynaptic neurons^39^, suggesting that UNC-7 is required to function in the mature AFD neuron.

### Cysteine mutants of UNC-7 failed to localize to the AFD axonal region

Innexins form two types of channels, gap junction channels and undocked hemichannels^40–42^. To address whether UNC-7 acts as a gap junction channel or a hemichannel for the regulation of thermotaxis, we utilized the cysteine mutants of UNC-7, UNC-7(Cysless; C173A, C191A, C377A, C394A) and UNC-7(C191A). These cysteine residues were previously reported to be essential for the formation of gap junctions but were dispensable for hemichannel activity^43^. We therefore asked whether the expression of *unc-7(Cysless)* or *unc-7(C191A)* in AFD could rescue the thermotaxis defect of *unc-7(AFD KO).* However, we observed that these mutants of UNC-7 failed to localize to the axonal region of AFD unlike the wild-type UNC-7, and that the expression of these mutant *unc-7* genes did not rescue the thermotaxis defect of *unc-7(AFD KO)* (Supplementary Fig. 2). Given that the cysteine mutants of UNC-7 localized in the neuronal processes of the ventral nerve cord^43^, these observations suggested that the cysteine residues are required for the transport of UNC-7 in the AFD sensory neurons and that UNC-7 might be transported by distinct mechanisms depending on the neuronal subtypes.

### UNC-7 acts as a hemichannel to regulate thermotaxis

Since the cysteine mutants of UNC-7 could not be utilized to address the type of channel that UNC-7 funcions during thermotaxis, we attempted to design a new form of UNC-7 that would lose the ability to form gap junctions but retain the hemichannel activity. Since previous structural studies suggested that the formation of a gap junction is mediated through the interaction between the second extracellular loops (EL2) of the opposing hemichannels^44^, we generated a chimeric form of UNC-7 whose coding sequence for the EL2 was replaced by that of mouse pannexin 1 (mPANX1), which belongs to the family of pannexin, the functional homolog of innexins^45–47^ (Fig. 2a). The AFD-specific expression of this chimeric UNC-7 rescued the thermotaxis defect of *unc-7(AFD KO)* animals, indicating that the activity of the chimeric UNC-7 is sufficient for regulating the thermotaxis (Fig. 2b). By contrast, expression of a full-length mPANX1 specifically in AFD did not rescue the thermotaxis defect of *unc-7(AFD KO)* (Supplementary Fig. 3a).

**Figure 2.**
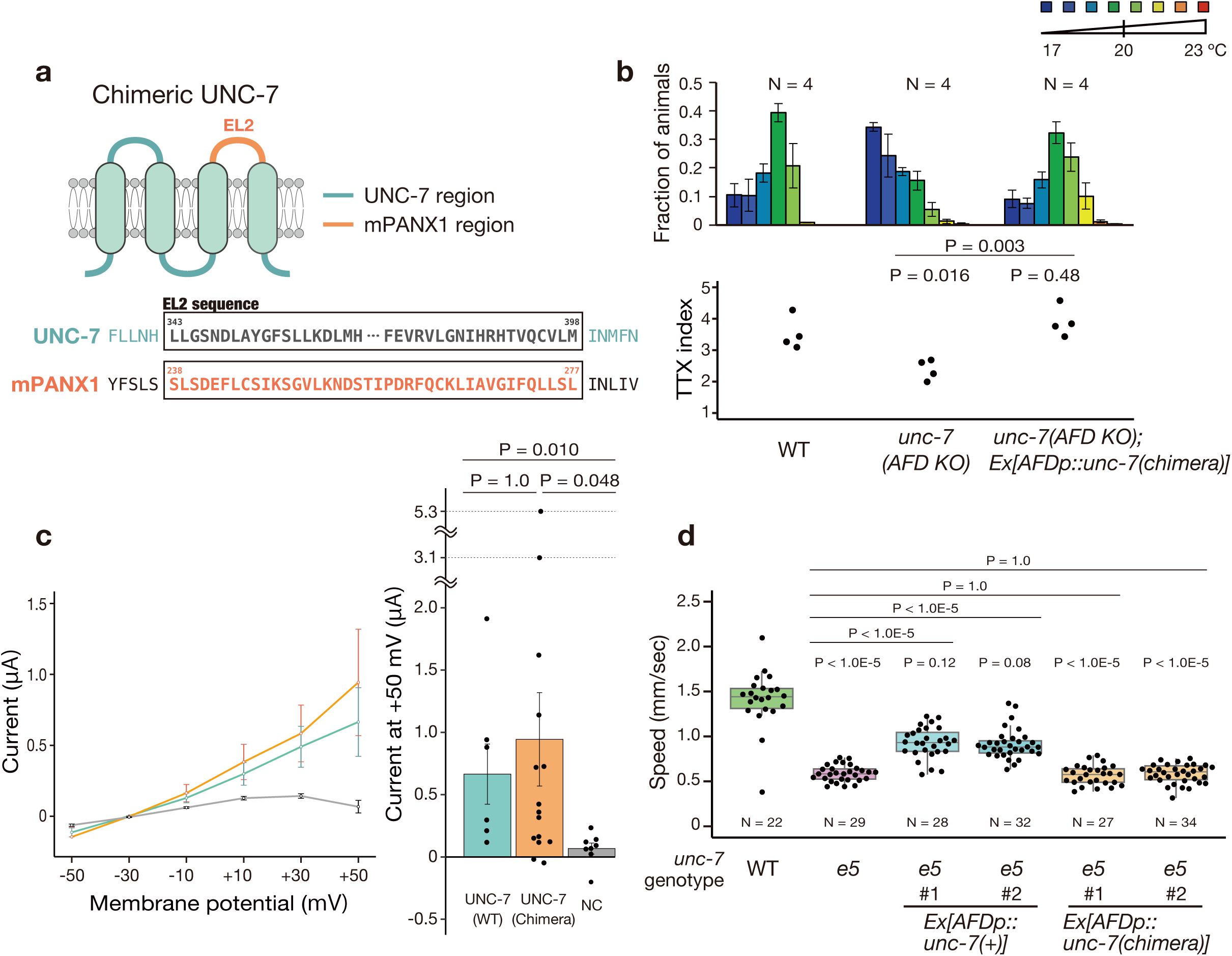
UNC-7 functions as a hemichannel to control the thermotaxis behavior. (a) The design of chimeric UNC-7 is shown. The sequence coding the second extracellular loop (EL2) of UNC-7 was replaced with that of Mouse Pannexin1 (mPANX1). (b) TTX behavior of *unc-7(AFD KO)* animals carrying a transgene that expresses the chimeric UNC-7 specifically in the AFD neuron. Distributions of the animals on the thermal gradients of the TTX plate are displayed in the top histograms. The fraction of the population in each temperature section is shown as mean ± SEM. TTX indexes are shown in the bottom dot plots. (c) Electrophysiological analysis using Xenopus oocyte to test hemichannel activity of the wild-type and chimeric UNC-7. Measured voltage clamp currents of the wild-type UNC-7, chimeric UNC-7 and negative control (NC) are indicated in the I/V plot as means ± SEMs (Left). The bar graphs indicate the means of currents at + 50 mV (Right). Individual data points are shown as dots. Error bars correspond to SEM. (d) Locomotion assay to test the gap junctional activity of chimeric UNC-7. The speeds of individual animals are shown as dots. Box indicates the first and third quartiles, and whiskers extend to 1.5 times the interquartile range. P-values were determined by 1-way ANOVA with Tukey HSD test in (b) or by Exact Wilcoxon rank sum test with Bonferroni correction in (c) and (d).

To assess whether the chimeric UNC-7 retains hemichannel activity, we expressed the chimeric UNC-7 in *Xenopus* oocyte and confirmed that the chimeric UNC-7 exhibited voltage-dependent hemichannel currents comparable to that observed by the wild-type UNC-7 (Fig. 2c, Supplementary Fig. 4). We also tested whether the chimeric UNC-7 displays the gap junction activity. It has been previsouly reported that the gap junction activity of UNC-7 is required in the motor neurons to mediate locomotion^48^. We therefore examined whether the chimeric UNC-7 can rescue the uncoordinated movement of animals carrying *unc-7(e5)* and found that the pan-neuronal expression of the wild-type UNC-7 partially rescued the locomotory defect of *unc-7(e5),* whereas the chimeric UNC-7 did not (Fig. 2d). These results indicated that the chimeric UNC-7 loses the ability to form gap junctions but retains the hemichannel activity. In addition, we observed that UNC-7(M121L), a mutant form of *unc-7* previously shown to be unable to form the gap junction^48^, also rescued the thermotaxis defect of *unc-7(AFD KO)* animals (Supplementary Fig. 3b). Together, these results suggested that the hemichannel activity of UNC-7 in AFD is sufficient to regulate thermotaxis.

### UNC-7 functions downstream of calcium influx in AFD

A previous report indicated that the UNC-7 hemichannel is required for the sensory transduction of mechanosensation in *C. elegans*^49^. Given this report, we assessed whether UNC-7 regulates temperature sensing in the AFD thermosensory neurons. The AFD neurons were previously shown to increase its calcium concentration in response to temperature warming, and this calcium response occurs around the cultivation temperature^22–26^. We therefore monitored calcium dynamics of AFD in wild-type, *unc-7(e5)* and *unc-7(AFD KO)* animals and found that the calcium response of AFD was normal in *unc-7* mutants (Fig. 3a-d). We did not detect significant differences in the maximum value of ratio change or response temperature of AFD between wild-type animals and *unc-7* mutants (Fig. 3c, d). These results indicated that *unc-7* does not affect temperature-evoked calcium response in AFD and suggested that UNC-7 regulates a process downstream of calcium influx in AFD.

**Figure 3.**
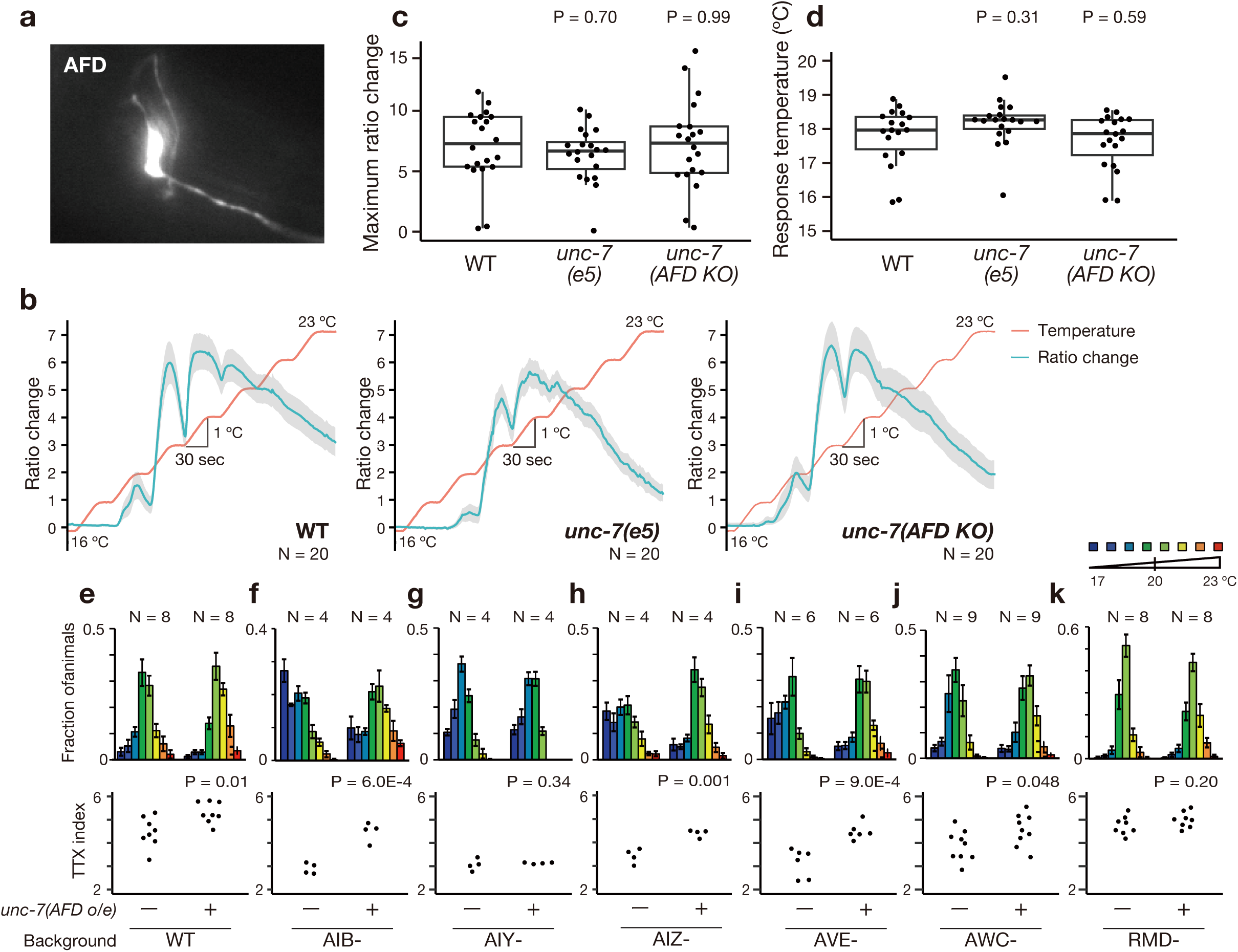
UNC-7 transmits neuronal information from the AFD thermosensory neurons to the AIY interneurons. (a) A fluorescent image of the AFD neuron expressing the GCaMP6m calcium indicator. (b) Ratio changes of the AFD calcium dynamics are shown. Blue lines indicate mean values and grey regions SEMs. N = 20 for each. (c) The maximum ratio changes in the wild type*, unc-7(e5)* and *unc-7(AFD KO)* animals are shown. P-values were determined by Dunnet test. (d) The response temperature of the AFD calcium response was defined as the temperature at which the ratio change first exceeded 0.5. P-values were determined by Steel test. (e-k) TTX behaviors of the wild-type and cell-ablated animals carrying a transgene that expresses the wild-type UNC-7 specifically in AFD. Distributions of the animals on the thermal gradients of the TTX plate are displayed in the top histograms. The fraction of the population in each temperature section is shown as mean ± SEM. TTX indexes are shown in the bottom dot plots. P-values were determined by Student’s T test in (e), (f), (h), (i), (j) and (k) or by Exact Wilcoxon rank sum test in (g).

### UNC-7 transmits temperature information to the AIY interneuron

To identify neurons to which UNC-7 hemichannels transmit temperature information from AFD, we conducted single cell ablation experiments by using reconstituted caspases^17,50^. We individually ablated a subset of the neurons reported to be chemical or electrical synapse partners of AFD^19^ and asked which neurons, when ablated, could suppress the effect of the AFD-specific overexpression of UNC-7. We observed that while overexpression of *unc-7* affected thermotaxis behaviors of animals lacking AIB, AIZ, AVE or AWC, the ablation of the AIY interneuron or the RMD motor neuron suppressed the defect caused by the *unc-7* overexpression (Fig. 3e-k). These results suggested that UNC-7 transmits temperature information to the AIY and RMD neurons. Since the AIY interneurons have been shown to play pivotal roles in thermotaxis^15,17,51^, we focused on the neural pathway between AFD and AIY and further investigated the mechanisms of the behavioral regulation by UNC-7 hemichannels.

### UNC-7 regulates curving bias by transmitting temperature information to AIY

Previous studies showed that multiple behavioral components such as turns, reversals and curves, are important for the regulations of thermotaxis behavior. Many of these behavioral components are regulated by the AFD-AIY neural pathway, and each of the behavioral components contributes to thermotaxis to a different degree^16,17^. To address which behavioral components are regulated by UNC-7, we performed high-resolution behavioral analysis using Multi Worm Tracking system^17,30,52^ (Fig. 4, Supplementary Fig. 5, 6). We cultivated animals at 20 °C and monitored their behaviors within the temperature range from 18.5 °C to 21.5 °C (Fig. 4a). Consistent with previous reports^17,30^, we observed that wild-type animals displayed curving bias toward warmer temperature when moving down the thermal gradient and curved toward colder temperature when moving up the thermal gradient. These bidirectional curving bias presumably drives the animals toward the cultivation temperature (Fig. 4). *unc-7(AFD KO)* animals showed defects in the regulation of curve and, in particular, displayed lower curving bias than that of the wild-type animals when moving down the thermal gradient (Fig. 4d). This curving defect of *unc-7(AFD KO)* was rescued by expressing the wild-type or the chimeric UNC-7 in AFD. These results suggested that UNC-7 hemichannels promote the curving bias toward warmer temperature when animals are moving down the temperature gradient away from the cultivation temperature, thereby promoting migration toward the cultivation temperature. We did not observe significant defects in other behavioral components examined, including omega turn, reversal, reversal turn, shallow turn or speed (Supplementary Fig. 5).

**Figure 4.**
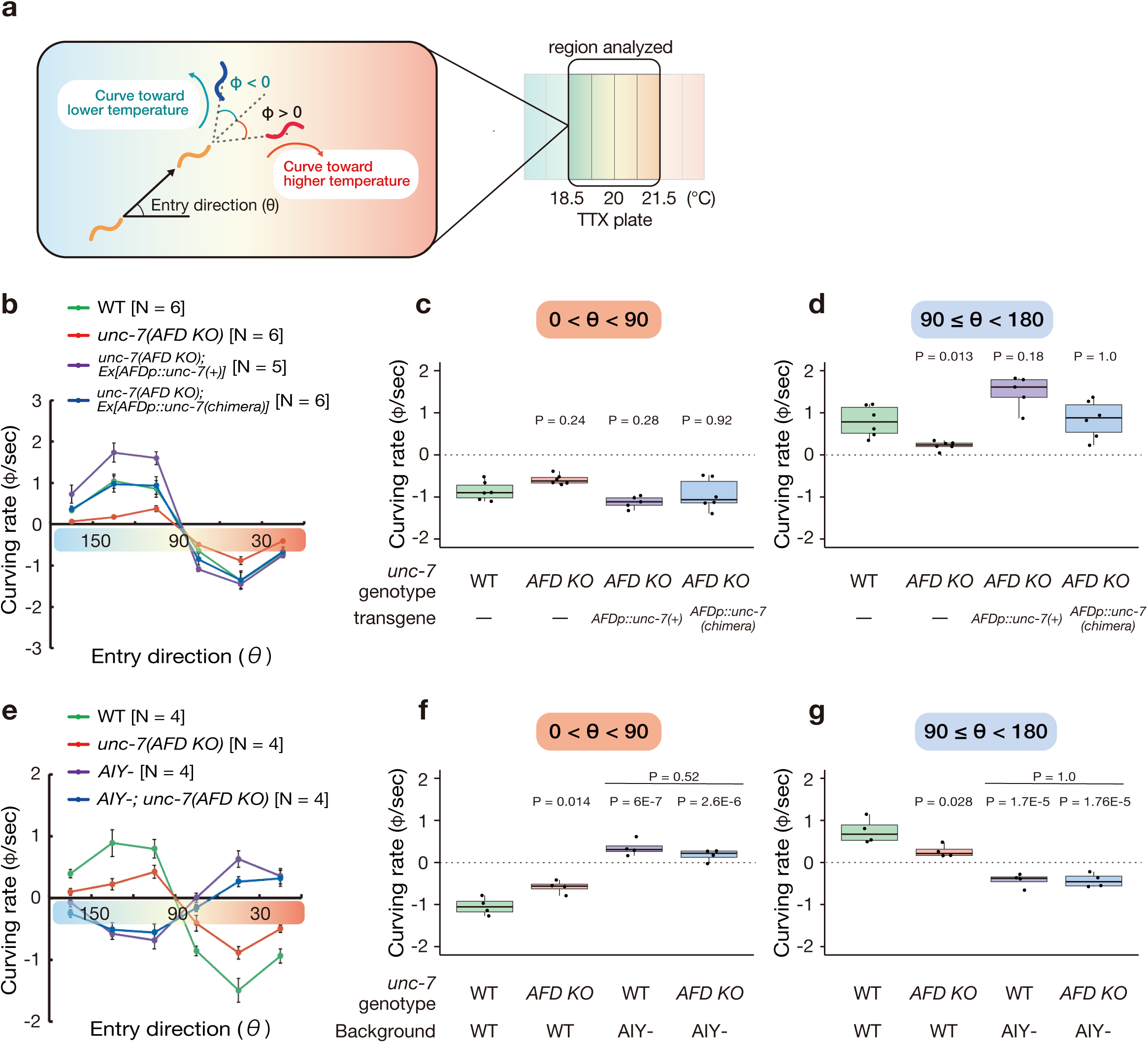
UNC-7 regulates the curving bias during thermotaxis. (a) Schematic of the analysis of the curve is shown. The entry direction θ is defined as the angle between the animal’s moving direction and the vector pointing to the warm side of the thermal gradient. The curving bias is measured as the angle (φ) between the past and current moving directions. We define φ as a positive value if animals curve toward the warm side and as a negative value when they curve toward the cold side. Behaviors of animals within the temperature range from 18.5 °C to 21.5 °C during 40 min from the start of the assay were analyzed. The cultivation temperature was 20 °C. (b) The Curving biases of the wild-type animals (green), *unc-7(AFD KO)* animals (red) and *unc-7(AFD KO)* animals carrying a transgene that expresses the wild-type (purple) or the chimeric UNC-7 (blue) specifically in the AFD thermosensory neuron are shown. The dots represent the means of curving rates of 50 - 400 animals from individual recordings. Error bars indicate SEM. (c) The curving biases of the animals moving up the thermal gradient (0 < θ < 90) are shown. The averages of the curving bias per individual recording are shown as dots. Box indicates the first and third quartiles, and whiskers extend to 1.5 times the interquartile range. P-values were determined by 1-way ANOVA with Tukey HSD test. (d) The Curving biases of the animals moving down the thermal gradient (90 =< θ < 180) are shown. P-values were determined by Exact Wilcoxon rank sum test with Bonferroni correction. (e-g) The Curving biases of the wild-type animals (green), *unc-7(AFD KO)* animals (red), AIY interneuron-ablated animals (purple) and AIY interneuron-ablated animals lacking *unc-7* specifically in the AFD neuron (blue) are shown as in (b-d). P-values were determined by 1-way ANOVA with Tukey HSD test in (f) and (g).

To ask whether this regulation of the curving bias by UNC-7 involves the AIY interneuron, we analyzed behavioral components of the AIY-ablated animals (Fig. 4e-g, Supplementary Fig. 7). We found that the AIY-ablated animals showed the opposite curving bias to that observed in the wild type: they curve toward colder temperature when moving down the thermal gradient and toward warmer temperature when moving up the thermal gradient (Fig. 4e-g). Importantly, *unc-7(AFD KO)* did not affect the curving bias in the AIY-ablated animals. Together, these results suggested that UNC-7 regulates curving bias while animals are moving toward colder temperature and that UNC-7 mediates transmission of temperature information from AFD to AIY.

### UNC-7 hemichannels inhibit the activity of the AIY neuron upon cooling stimuli

To assess whether UNC-7 regulates the neuronal activity of the AIY interneuron, we analyzed calcium dynamics of AFD and AIY in freely behaving animals using the microscope with an automated tracking system (Fig. 5). Since our behavioral component analysis revealed that UNC-7 regulates curve when animals were migrating down the temperature gradient, we subjected the animals to a cooling stimulus, where the temperature was decreasing away from the cultivation temperature: 20 °C (Fig. 5b). We observed that in response to this cooling stimulus, the wild-type AFD neurons showed decreases in the calcium concentration (Fig. 5c). This observation is consistent with the previous report that cooling hyperpolarizes the AFD neurons^24^. We also found that the majority of the AIY neurons of the wild-type animals responded to the cooling stimulus by decreasing the calcium concentration (Fig. 5c). This result corresponds to our previous report that the AIY responses correlate with the valence of thermal stimuli, with stimuli with positive valence evoking excitatory responses and stimuli with negative valence inhibitory responses^30^. When *unc-7(AFD KO)* animals were subjected to cooling stimuli, the activity of AFD was decreased similarly to those observed in the wild-type animals. By contrast, decreases in the AIY calcium concentration were attenuated in *unc-7(AFD KO)* animals when compared to those in the wild-type animals. To compare the AIY responses between the wild-type and *unc-7(AFD KO)* animals, we analyzed the proportion of standardized fluorescent change below -0.3 and found that the ratio in *unc-7(AFD KO)* is lower than that in the wild type (Fig. 5d). This defect was rescued by AFD-specific expression of the wild-type UNC-7 (Fig. 5c, d) or of the chimeric UNC-7 (Fig. 5e, f). We also monitored the neuronal activities in animals subjected to warming stimuli and found that *unc-7(AFD KO)* did not affect the regulation of the AIY activity upon warming (Fig. 5g-i). These results indicated that the hyperpolarizing AFD neuron inhibits the activity of the AIY neuron in response to cooling stimuli and that UNC-7 hemichannels regulate this process.

**Figure 5.**
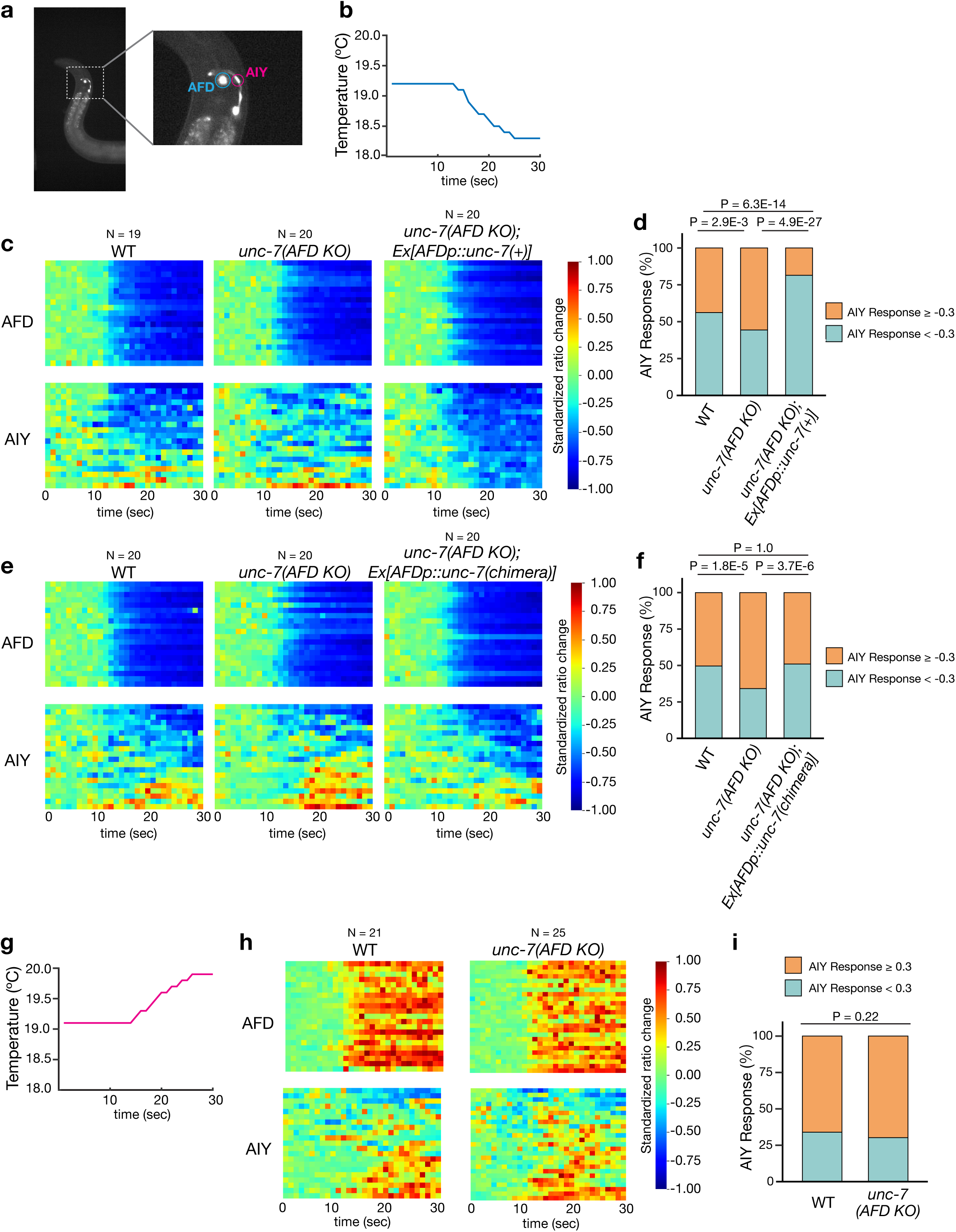
UNC-7 functions to suppress the AIY interneuron in response to the cooling stimulus. (a) A fluorescent image of an animal expressing the YCX calcium indicator in the nucleus of the AFD neuron (blue circle) and AIY neuron (red circle). (b) A representative of the temperature program for cooling is shown. (c) Calcium imaging of AFD and AIY neurons in the wild-type, *unc-7(AFD KO)* and *unc-7(AFD KO)* animals carrying a transgene that expresses the wild-type UNC-7 in response to cooling stimulus. Standardized fluorescence ratio changes of AFD (top) and AIY (bottom) are displayed in the heat maps. Each row corresponds to individual recording. (d) The percentage of the AIY data frame in which the standardized ratio change is over -0.3 or below -0.3 is shown in the bar graph. P-values were determined by Chi-Square test with Bonferroni correction. (e) Calcium imaging of AFD and AIY neurons in the wild-type, *unc-7(AFD KO)* and *unc-7(AFD KO)* animals carrying a transgene that expresses the chimeric UNC-7 in response to cooling stimulus. Standardized fluorescence ratio changes are shown as in (c). (F) The percentage of AIY response per category is shown as in (d). (g) A representative of the temperature program for warming is shown. (h) Calcium imaging of AFD and AIY neurons in the wild-type and *unc-7(AFD KO)* animals in response to warming stimulus. Standardized fluorescence ratio changes are shown as in (c). (i) The percentage of the AIY data frame in which the standardized ratio change is over 0.3 or below 0.3 is shown in the bar graph. P-values were determined by Chi-Square test with Bonferroni correction.

### UNC-7 regulates thermotaxis independently of synaptic vesicle exocytosis

We next address whether inhibition of the AIY activity by UNC-7 is mediated by controlling the synaptic transmission from AFD. The AFD-AIY synaptic transmission was shown to be brought about by two kinds of the neurotransmitters, neuropeptides for excitatory signaling and glutamates for inhibitory signaling^27,28^. The glutamatergic transmission from AFD required *eat-4*, which encodes a vesicular glutamate transporter (VGLUT) that transports glutamates into synaptic vesicles^28,53^. To assess whether UNC-functions through controlling synaptic release of glutamates, we conducted the epistasis analysis between *unc-7* and *eat-4*. We generated animals lacking *eat-4* specifically in AFD by the Cre/*loxP* system and found that *eat-4(AFD KO)* animals displayed thermophilic defects (Fig. 6a), indicating that EAT-4 acts in AFD to regulate thermotaxis behavior. Importantly, the *unc-7(AFD KO)* mutation affected thermotaxis in animals lacking *eat-4* in AFD. These results indicated that UNC-7 acts in parallel to EAT-4 to regulate thermotaxis.

**Figure 6.**
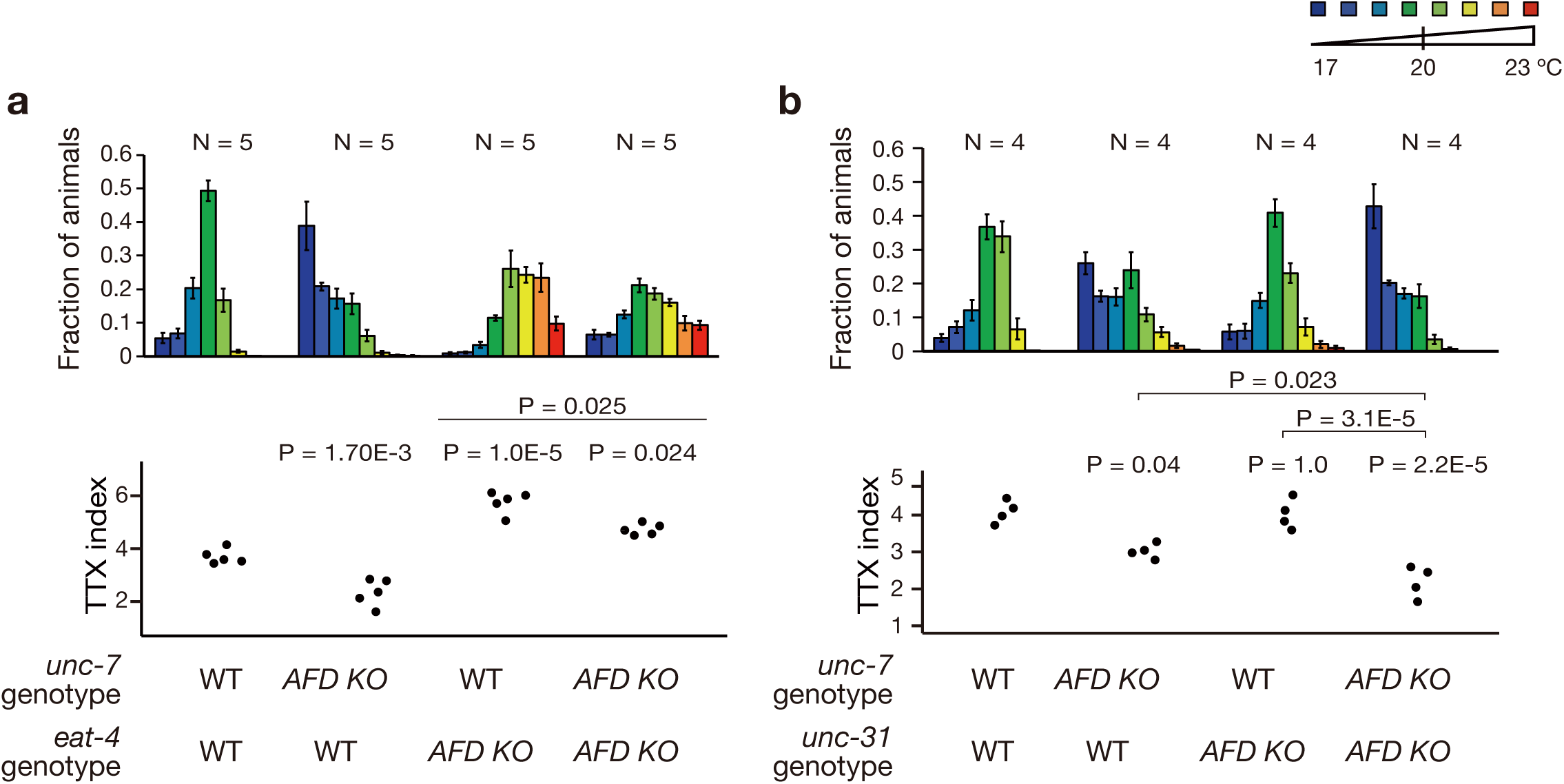
*unc-7(AFD KO)* affected the thermotaxis in animals lacking *eat-4* or *unc-31* in the AFD thermosensory neuron. (a) TTX behaviors of animals lacking *unc-7* and *eat-4* specifically in the AFD thermosensory neuron. (b) TTX behavior of animals lacking *unc-7* and *unc-31* specifically in the AFD thermosensory neuron. Distributions of the animals on the thermal gradients of the TTX plate are displayed in the top histograms. The fraction of the population in each temperature section is shown as mean ± SEM. TTX indexes are shown in the bottom dot plots. P-values were determined by 1-way ANOVA with Tukey HSD test.

We also tested the possibility that UNC-7 is involved in the peptidergic signaling within AFD. Previous studies indicated that *unc-31*, which encodes the calcium dependent activator protein essential for secreting neuropeptides, is required for the peptidergic signaling between AFD and AIY^27,54^. We tested the epistasis between *unc-7* and *unc-31* and found that while *unc-31(AFD KO)* did not display a thermotaxis defect, animals lacking both *unc-7* and *unc-31* showed a cryophilic phenotype stronger than that of *unc-7(AFD KO)* or *unc-31(AFD KO)* animals (Fig. 6b). These results suggested that UNC-31 plays at least a minor role in AFD to regulate thermotaxis and that UNC-7 acts in parallel to UNC-31. Furthermore, we examined thermotaxis behaviors of single mutants for a panel of the neuropeptide genes that are expressed in AFD^55,56^ and found that none of the mutants tested displayed a thermotaxis defect (Supplementary Fig. 7). These results implied that UNC-7 functions to regulate thermotaxis independently of the synaptic vesicle exocytosis.

## Discussion

In this study, we report that the AFD sensory neurons inhibit the activity of the AIY interneurons while animals are subjected to cooling stimuli and that UNC-7 hemichannels function in AFD to regulate this process. Since our imaging results and the previous study indicate that cooling stimuli hyperpolarize the AFD neuron^24^, these results suggest that the hyperpolarizing AFD neurons recruit UNC-7 hemichannels to downregulate the AIY activity.

Our results uncovered the roles of innexin hemichannels in promoting neurotransmission from hyperpolarized cells. Previously, gap junctions have been shown to mediate the propagation of hyperpolarization from one cell to the next. For example, in mammals, hyperpolarizing current spreads directly from the endothelial cells to the smooth muscle cells via myoendothelial gap junctions composed of connexins^57,58^. In mammalian cerebellum, gap junctions composed of connexins between Golgi neurons were shown to facilitate the propagation of hyperpolarization after a brief depolarizing current^12,13^. In *C. elegans,* the gap junctions composed of UNC-9, a member of innexin family proteins, were also implicated to spread membrane hyperpolarization from RIM motor neurons to AVA interneurons during spontaneous locomotor activity^59^. These reports suggest that the gap junctions composed of innexins and connexins mediate the propagation of membrane hyperpolarization. Our findings now indicate that innexins can also function as hemichannels to regulate neurotransmission from hyperpolarizing neurons. We suggest that hemichannel-composing proteins -innexins, connexins and pannexins-could be general mediators of neural transmission in hyperpolarized cells.

Recent studies in mammals suggested that glial cells employ hemichannels to modulate communications between neurons. For example, the connexin hemichannels expressed in astrocytes contribute to excitatory synaptic transmission from CA3 to CA1 pyramidal neurons by releasing ATP and glutamine^8,9^. Another report suggested that astroglial hemichannels release chemokines to enhance spinal cord synaptic transmission^60^. Thus, these observations led to the notion that hemichannels in glial cells support neurotransmission. Our findings further advance this idea and show that neuronal UNC-7 hemichannels directly regulate neural communication. We speculate that neuronal hemichannels composed of innexins and possibly those of connexins and pannexins could similarly contribute to neurotransmission.

Our epistasis analysis suggested that UNC-7 hemichannels expressed in the AFD neurons regulate neurotransmission independently of chemical synaptic transmissions. The most plausible function of UNC-7 would be to release transmitters. In mammals, channel proteins had been reported to release neurotransmitters to signal sensory information: the CALHM1/CALHM3 channels are essential for taste perception and mediate depolarization-dependent release of ATP from taste cells^61^. Like CALHM1/CALHM3 channels, UNC-7 hemichannels might allow neurotransmitters to pass through, and such neurotransmitters exert inhibitory effects on the AIY neuronal activity. This releasing mechanism by hemichannels is likely distinct from that by CALHM channels in that it would operate in the absence of calcium influx. Given that innexins form a large pore channels^42^, innexin hemichannels can facilitate the passage of diverse ligands. Depending on a combination of ligands and their cognate receptors, hemichannel-mediated neurotransmission might allow diverse modes of neural communication from hyperpolarized cells. Our results thus suggest a previously unrecognized mechanism of neurotransmission that expands the way of the neural circuitry operations.

How does this mechanism of the innexin-mediated neurotransmission contribute to thermotaxis behavior? The AIY neurons regulate the curving bias as well as the frequency of reversals and turns^17,62^, and the reduction in the AIY activity generally correlates with reorientation maneuvers^16,63,64^. Thus, the AIY activity is expected to be inhibited when animals are moving away from the cultivation temperature on a thermal gradient. Consistent with this expectation, previous studies indicated that when animals migrate up a thermal gradient above the cultivation temperature, AFD depolarizes upon warming stimulus, triggers glutamate release via chemical synapse and inhibits AIY activity^28,30^. The AFD neurons were also reported to play an essential role in behavioral control while animals were moving down a thermal gradient below the cultivation temperature^17^. However, since the AFD membrane would be hyperpolarized under such thermal context, how AFD controls the neural circuitry had remained unknown. Our studies provide an answer to this question and reveal that UNC-7 hemichannels function as calcium-independent regulators of neurotransmission and inhibit the AIY neurons. We speculate that such neurotransmissions via hemichannels in response to membrane hyperpolarization can play crucial roles in various sensory contexts. The careful investigation of the regulatory mechanism by the innexin hemichannels will lead to a further understanding of neurotransmission in hyperpolarizing neurons.

## Methods

### C. elegans strains

*C. elegans* animals were cultivated at 20 °C on nematode growth media (NGM) plates seeded with *E. coli* OP50 bacteria^32^. N2 (Bristol) was used as the wild-type strain. Germ line transformation was performed by microinjection as previously described^65^. Genome editing was performed by the CRISPR Cas9 system as previously described^66^. All strains used in this study are shown in Supplementary Table 1.

### Thermotaxis assay

Thermotaxis (TTX) assays were performed as previously described^67^. A linear thermal gradient from 17 °C to 23 °C was formed on a thermotaxis assay plate with a temperature steepness of approximately 0.5 °C/cm. The center of the plate was set at 20 °C. Animals cultivated at 20 °C were placed on the center of a thermotaxis assay plate and were allowed to freely behave for 1 hour. We divided the thermotaxis assay plate into sections along the temperature gradient, and the number of animals in each section was counted. We calculated thermotaxis (TTX) indexes using the formula shown in Figure 1a.

### Locomotion rescue experiment

We tracked and recorded the behaviors of ∼30 animals on assay plates for 5 min by using Multi Worm Tracking system^52^. Instantaneous speeds of individual animals tracked for longer than 2 consecutive minutes were calculated by Chreography^52^, and the average speeds of individual animals were calculated.

### Electrophysiological analysis

Whole-cell voltage clamp recording of Xenopus oocytes was performed as previously described^68^. The wild-type and chimeric *unc-7* genes were cloned into pGEM-HeFx plasmids, and the cRNA was generated from each plasmid by using an RNA preparation kit (mMessage mMachine T7 Transcription Kit; Invitrogen) according to the manufacturer’s protocol. The oocytes were collected from *Xenopus laevis* and then treated with collagenase solution, which contains 2 mg/mL collagenase type I (Gibco) dissolved in OR2 buffer (82.5 mM NaCl, 2 mM KCl, 1 mM MgCl_2_, and 5 mM HEPES [adjusted to pH 7.5 with NaOH]) at 18 °C for 1.5 h. Forty nano grams of *unc-7* cRNA were coinjected with 10 ng of antisense oligonucleotide DNA for Xenopus Cx38 into oocytes. The oocytes for negative controls were injected with the antisense DNA only. The oocytes were incubated at 18 °C for 2∼3 days in ND96 buffer (93.5 mM NaCl, 2 mM KCl, 1.8 mM CaCl_2_, 2 mM MgCl_2_, and 5 mM HEPES [adjusted to pH 7.5 with NaOH]). Hemichannel currents were recorded by using iTEV 90 Multi-Electrode Clamp Amplifier (HEKA Elektronik). The oocytes placed in ND96 buffer without CaCl_2_ were clamped at -30 mV initially and then subjected to 10 sec voltage steps from -50 mV to +50 mV in 20 mV increments.

### Critical period analysis by AID system

We generated *unc-7(nj291)* strain, in which a degron sequence was inserted at the *unc-7* locus, and introduced into this strain a transgene that drives an AFD-specific expression of TIR1, an auxin-dependent E3 ubiquitin ligase. We added 1mM auxin indole-3-acetic acid (IAA) into NGM and TTX plates to allow TIR1 to bind degron-tagged UNC-7 and promote their ubiquitylation, leading to the degradation of degron-tagged UNC-7 by the proteasome specifically in AFD^38^.

### Calcium imaging of the AFD neurons in immobilized animals

Calcium imaging recordings of the AFD neurons in immobilized animals were performed as previously described^25^. Animals expressing calcium probe GCaMP6m together with TagRFP in AFD^69^ were cultivated at 20 °C and were immobilized on 10% agarose pads by polystyrene beads. We subjected animals to stepwise temperature warming and recorded fluorescence images of the AFD cell body by EM-CCD camera with 400 msec pulsed illumination every 1 sec. The fluorescence intensities were acquired from these images using MetaMorph software, and the ratio of fluorescence intensity (GCaMP3/TagRFP) was calculated. The ratio change was calculated by the following formula: (ratio – baseline ratio)/baseline ratio, where the baseline ratio value was the mean of the ratio values during the first 10 sec before the beginning of the temperature change.

### Calcium imaging of the AFD neurons and the AIY neurons in freely behaving animals

Calcium imaging of the AFD neurons and the AIY neurons in freely behaving animals were performed as previously described^30,70^. Animals expressing calcium probe YCX simultaneously in the AFD nuclei and the AIY neurons were cultivated at 20 °C and allowed to freely behave on 2 % agarose pads. We subjected animals to warming or cooling stimuli and tracked their movement. We recorded fluorescence images of AFD and AIY under the epifluorescent microscope with SOLA LED light engine as a light source with 30 msec pulsed illumination every 1 sec for 30 sec.

The fluorescence intensities from these images were analyzed using the MATLAB program^70^, and the ratio of fluorescence intensity (GCaMP3/TagRFP) was calculated. The intensities of AFD were acquired from its nuclei and that of AIY were acquired from a part of the axonal region shown in Fig. 5a. The ratio of fluorescence intensity (YFP/CFP) was standardized to 0-1 range, and the standardized ratio change was calculated by the following formula: (ratio – baseline ratio)/baseline ratio, where the baseline ratio value was the mean of the ratio values during first 10 sec before the beginning of the temperature change.

### Behavioral analysis

Multi Worm Tracking (MWT) assays were performed as previously described^17,30^. Animals cultivated at 20 °C were placed on the thermotaxis assay plate, and their behaviors were recorded using a CMOS camera (8 bits, 4,096 × 3,072 pixels; CSC12M25BMP19-01B, Toshiba-Teli) for 1 hour. MWT system extracted the coordinates of individual animal’s centroids and spines from the recording video. Using these data, the behavioral components of the worms on the temperature gradient within the range of 18.5 °C to 21.5 °C were analyzed by the MATLAB program previously described^17^.

### Statistics

Normality of the data was tested by Shapiro-Wilk test. Equality of variance among the data set was assessed by Bartlett test. When the normality and equality of variance of the data were assumed, Student t test was performed for paired comparison, and One-way analysis of variance (ANOVA) with Tukey–Kramer test or Dunnett test for multiple comparisons. When we could not assume the normality and equality of variance of the data, Exact Wilcoxon rank sum test was performed for paired comparison, and Exact Wilcoxon rank sum test with Bonferroni correction or Steel test were performed for multiple comparisons. Chi-Square tests with Bonferroni correction were performed for multiple comparisons of the percentage of a categorical AIY response in Fig. 6.

## Data availability

The data supporting the finding of this work are available within the paper and its Supplementary Information files. Source data are provided with this paper. Custom codes for the analysis of the calcium imaging experiments were previously published and are available at https://github.com/ShunjiNakano/AIY_tracking. The Multi-worm tracking data were analyzed by the custom codes previously published and are available at https://sourceforge.net/projects/mwt/files/ and https://github.com/ikedamuneki/ThermotaxisAnalysis.

## Acknowledgements

We thank K. Ikegami, Y. Murakami, and M. Murase for technical and administrative assistance; K. Noma for *knjIs16* strains; R. Ahluwalia for *nlp-50(nj340)* strains; A. Kano for help to rewrite the source code of behavioral analysis; the members of the Mori laboratory and Noma group at Nagoya University for discussions. Some strains were provided by *Caenorhabditis* Genetic Center, which is funded by NIH Office of Research Infrastructure Programs (P40 OD010440) or by NBRP, which is funded by the Japanese government. This work was supported by JST SPRING Grant Number JPMJSP2125 (to A.N.) and by JSPS KAKENHI Grant Numbers 23H02418 (to A.O.), 19H05644 (to I.M.), 18H05123 (to S.N.) and 21H052525 (to S.N.).

## Author contributions

A.N., I.M. and S.N. conceived the research. A.N., M.W., A.O., I.M. and S.N. designed the experiments. A.N., M.W., R.Y., H.K. and S.N. performed the experiments. A.N. and S.N. analyzed the data. A.N., I.M. and S.N. wrote the manuscript. All authors of this paper read and approved the final manuscript.

**Correspondence** and requests for materials should be addreassed to Shunji Nakano.

---

**Supplementary Figure 1.**
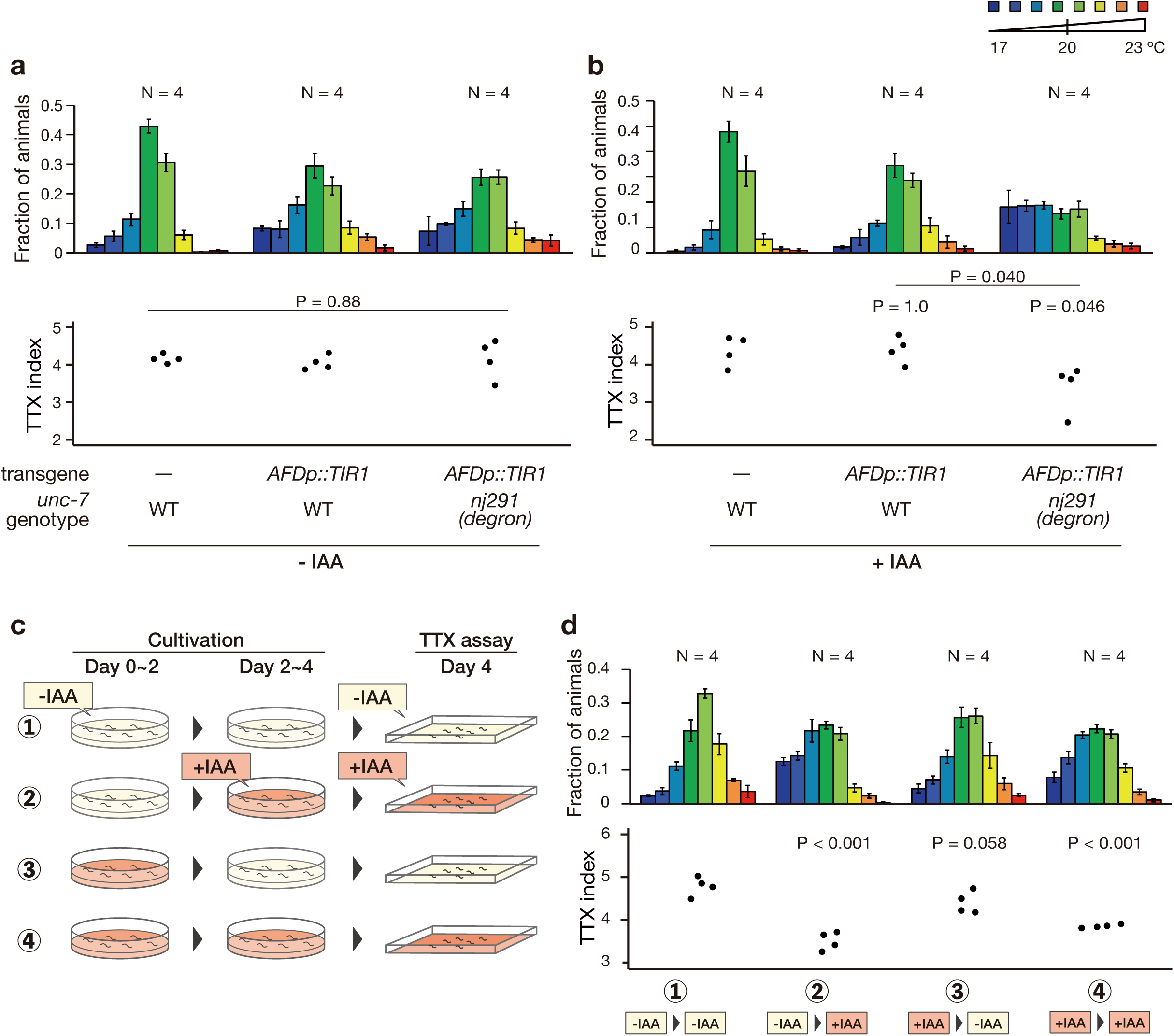
Critical time period analysis of UNC-7 by Auxin Induced Degradation system. (a, b) TTX behavior of *unc-7(nj291)* animals carrying a transgene that expresses TIR1, an auxin-dependent E3 ubiquitin ligase, specifically in the AFD thermosensory neuron. The degron sequence was inserted at the *unc-7* locus in *unc-7(nj291)* animals. The animals were grown and were subjected to thermotaxis assays in the absence (a) or the presence (b) of 1 mM auxin indole-3-acetic acid (IAA). Distributions of the animals on the thermal gradients of the TTX plate are displayed in the top histograms. The fraction of the population in each temperature section is shown as mean ± SEM. TTX indexes are shown in the bottom dot plots. P-values were determined by 1-way ANOVA with Tukey HSD test. (c) Schematic of critical time period analysis of UNC-7 by Auxin Induced Degron system. We cultivated animals in the presence or the absence of 1mM IAA for 4 days throughout their life - from embryo to adulthood- and examined their thermotaxis behaviors in the presence (④) of the absence (①) of IAA, respectively. We also subjected animals to a shift in the IAA condition from 0 to 1 mM (②) or from 1 to 0 mM (③) on day 2 when a majority of the animals were in their second or third larval stage. (d) TTX behaviors of *unc-7(nj291)* animals carrying a transgene that expresses TIR1 specifically in the AFD neuron. Animals were cultivated and subjected to thermotaxis assays under the conditions shown in (c) (① ∼ ④). P-values were determined by Dunnet test.

**Supplementary Figure 2.**
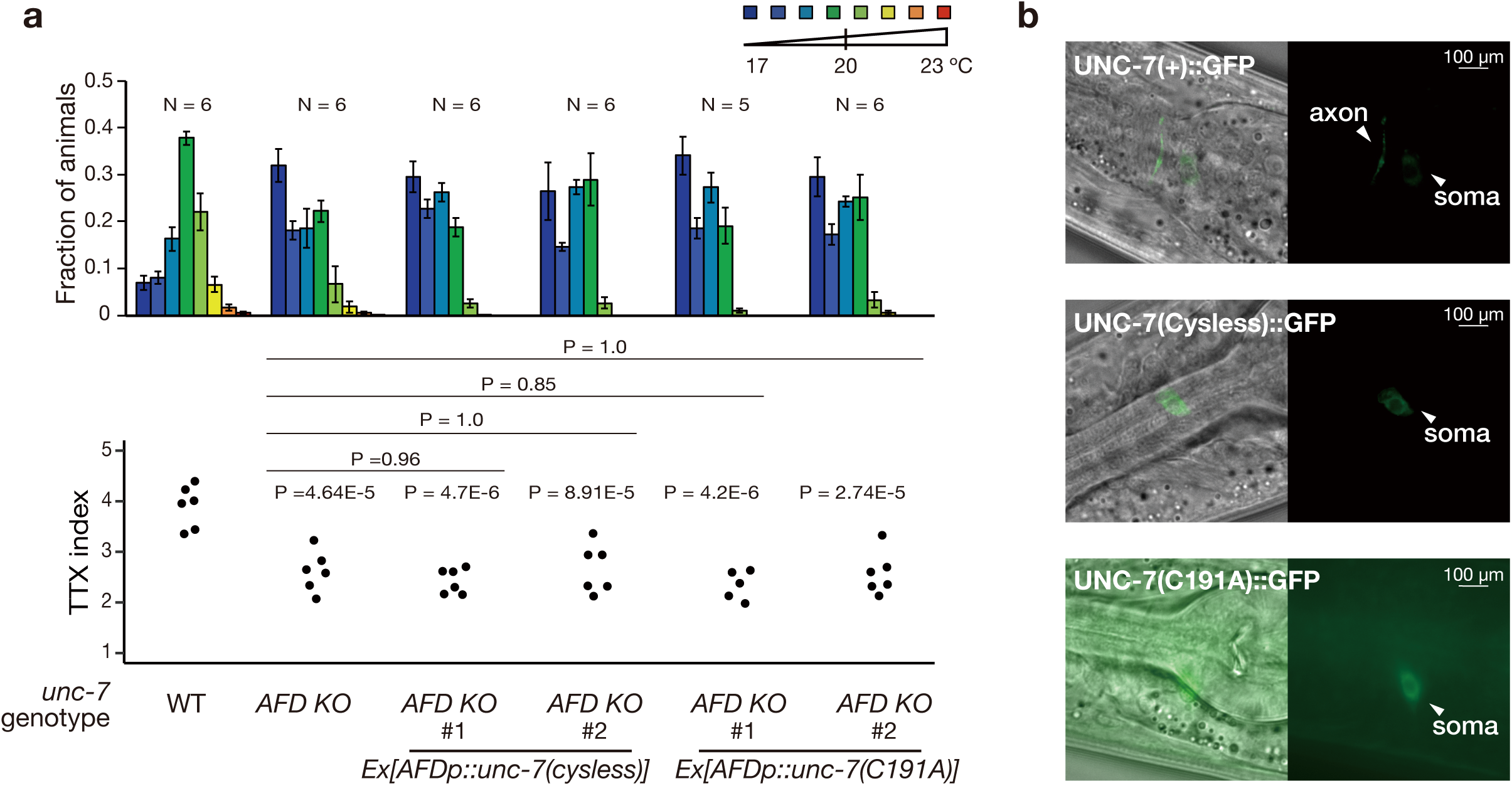
Cysteine mutants of UNC-7 fail to localize to the axonal regions of the AFD neuron. (a) TTX behavior of *unc-7(AFD KO)* animals carrying a transgene that expresses UNC-7(Cysless) or UNC-7(C191A) specifically in the AFD neuron. Distributions of the animals on the thermal gradients of the TTX plate are displayed in the top histograms. The fraction of the population in each temperature section is shown as mean ± SEM. TTX indexes are shown in the bottom dot plots. P-values were determined by 1-way ANOVA with Tukey HSD test. (b) Localization of the wild-type and cysteine mutants of UNC-7 in the AFD neuron. The wild-type or cysteine mutants of *unc-7* cDNA was fused with GFP, and each transgene was expressed under the *gcy-8L* promoter. Transmitted light images (Left) and fluorescent images (Right) are shown. The wild-type UNC-7::GFP localized at the soma and axon, whereas UNC-7(Cysless)::GFP and UNC-7(C191A)::GFP were localized exclusively at the soma of AFD.

**Supplementary Figure 3.**
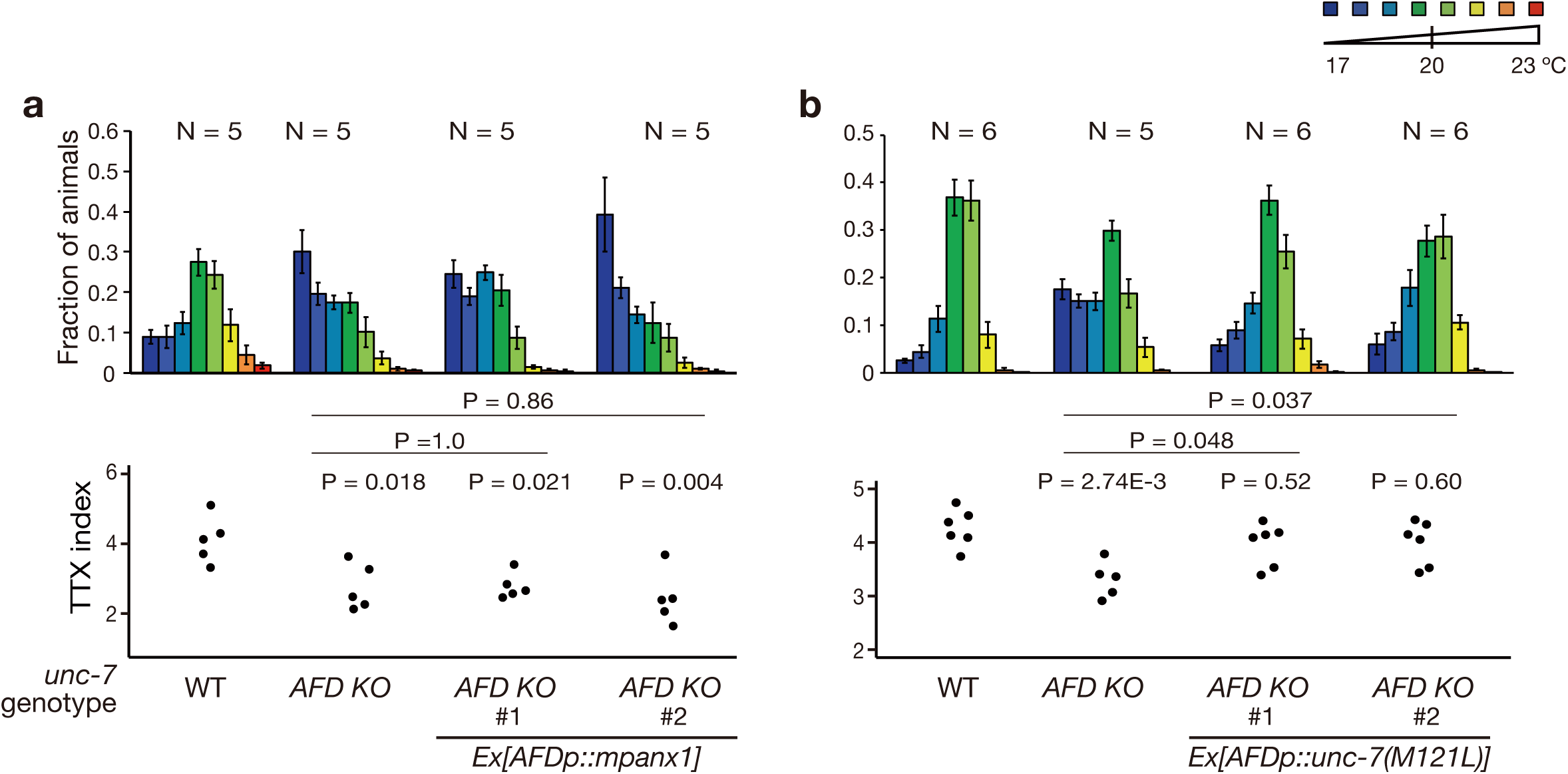
AFD-specific expression of UNC-7(M121L) rescued the thermotaxis defect of *unc-7(AFD KO)*. (a) TTX behavior of *unc-7(AFD KO)* animals carrying a transgene that expresses mPANX1 specifically in the AFD neuron. (b) TTX behavior of *unc-7(AFD KO)* animals carrying a transgene that expresses UNC-7(M121L) specifically in the AFD neuron. Distributions of the animals on the thermal gradients of the TTX plate are displayed in the top histograms. The fraction of the population in each temperature section is shown as mean ± SEM. TTX indexes are shown in the bottom dot plots. P-values were determined by 1-way ANOVA with Tukey HSD test in (a) and (b).

**Supplementary Figure 4.**
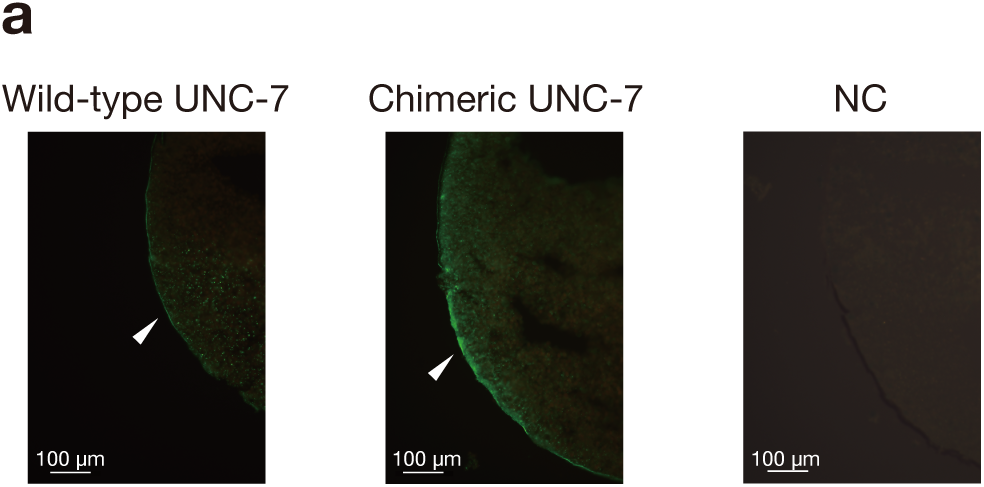
Expression and localization of the wild-type and chimeric UNC-7 in Xenopus oocytes. (a) Representative fluorescent images of Xenopus oocytes expressing the wild-type and chimeric form of UNC-7::GFP (green). NC: negative control of oocytes without UNC-7 injection. White arrows indicate UNC-7::GFP signals observed at the cell surfaces.

**Supplementary Figure 5.**
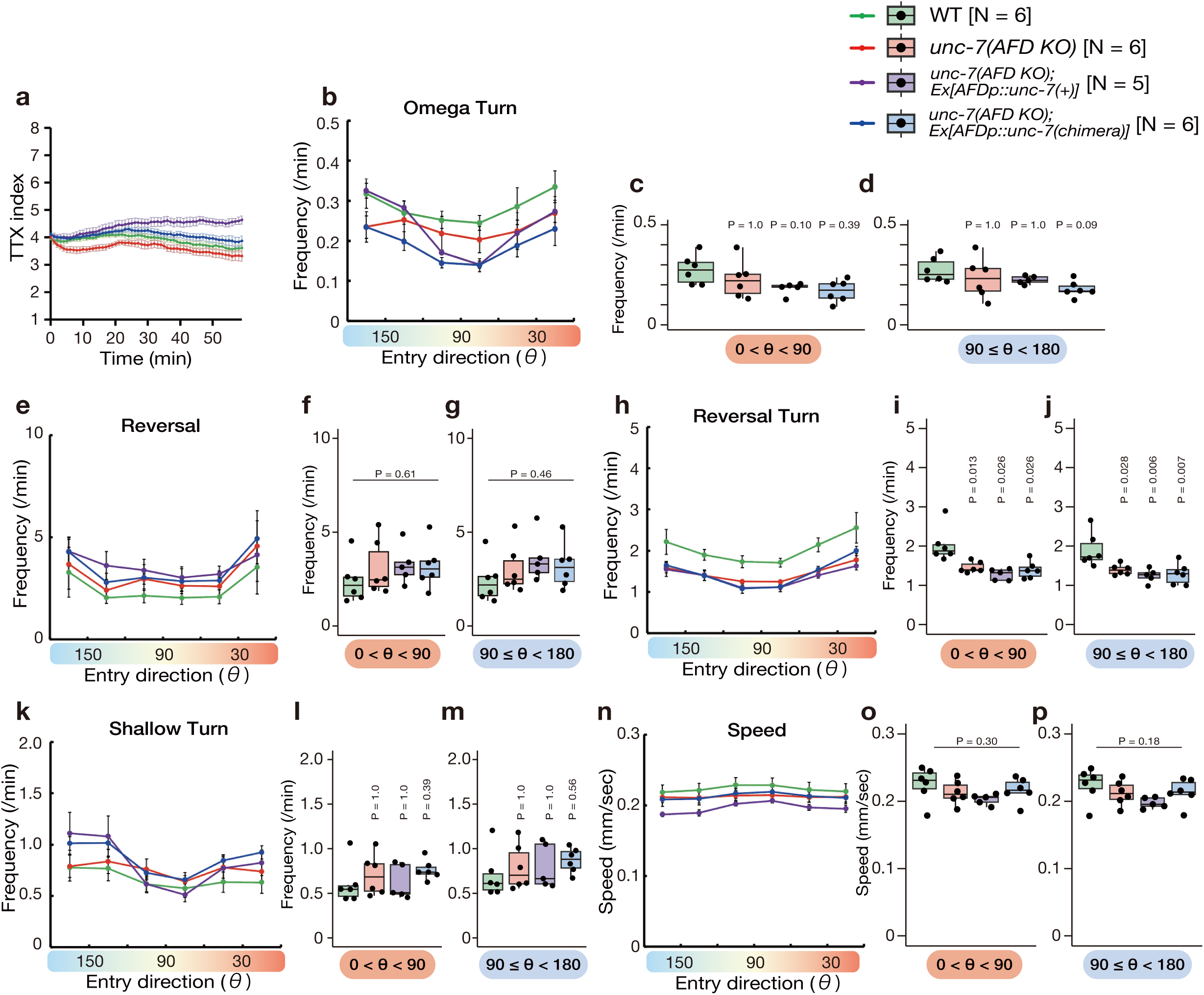
Behavioral component analysis of *unc-7* mutant animals. Behavioral component analysis of the wild-type animals (green), *unc-7(AFD KO)* animals (red) and *unc-7(AFD KO)* animals carrying a transgene that expresses the wild-type (purple) or the chimeric UNC-7 (blue) specifically in the AFD thermosensory neuron. The cultivation temperature was 20 °C. Animals behaving within the temperature region between 18.5 °C and 21.5 °C during the first 40 min of the recordings were analyzed for their behavioral components. Each recording is typically composed of 50-400 animals. We divided the animals’ entry directions into 30° bins and calculated the frequencies of each behavioral event when animals were moving in each direction. (a) The time course of TTX index. The dots and error bars indicate the means and SEMs. (b-m) The frequencies of omega turn (b-d), reversal (e-g), reversal turn (h-j) and shallow turn (k-m) are shown. The data in the line graphs are mean ± SEM. The frequencies of each behavioral component while the animals are moving up the thermal gradient [0 < θ < 90, (c), (f), (i) and (l)] or moving down the gradient [90 =< θ < 180, (d), (g), (j) and (m)] are shown. The averages of frequencies per individual recording are shown as dots. Box indicates the first and third quartiles, and whiskers extend to 1.5 times the interquartile range. (n-p) The averages of the speed of animals moving in each direction are shown as in (b-d). P-values were determined by Exact Wilcoxon rank sum test with Bonferroni correction in (c), (d), (i), (l) and (m), by 1-way ANOVA in (f), (g), (o) and (p) or by 1-way ANOVA with Tukey HSD test in (j).

**Supplementary Figure 6.**
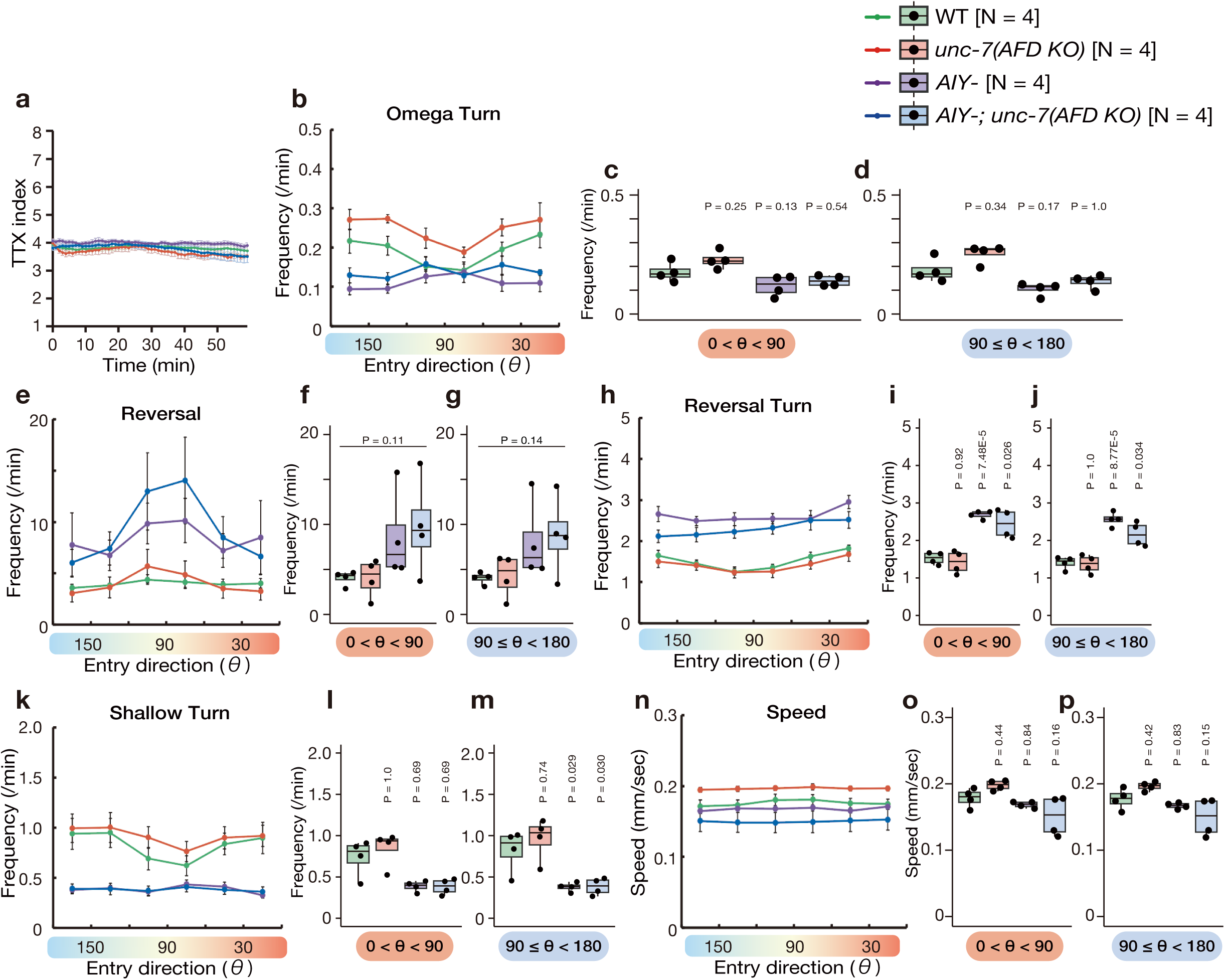
Behavioral component analysis of animals lacking the AIY neuron. (a-p) Behavioral component analysis of the wild-type animals (green), *unc-7(AFD KO)* animals (red), AIY interneuron-ablated animals (purple) and AIY interneuron-ablated animals lacking *unc-7* specifically in the AFD neuron (blue). The experiments were conducted, and the data are represented as in Supplementary Fig. 5. P-values were determined by 1-way ANOVA with Tukey HSD test in (c), (i), (j), (m), (o) and (p), by Exact Wilcoxon rank sum test with Bonferroni correction in (d) and (l) or by 1-way ANOVA in (f) and (g).

**Supplementary Figure 7.**
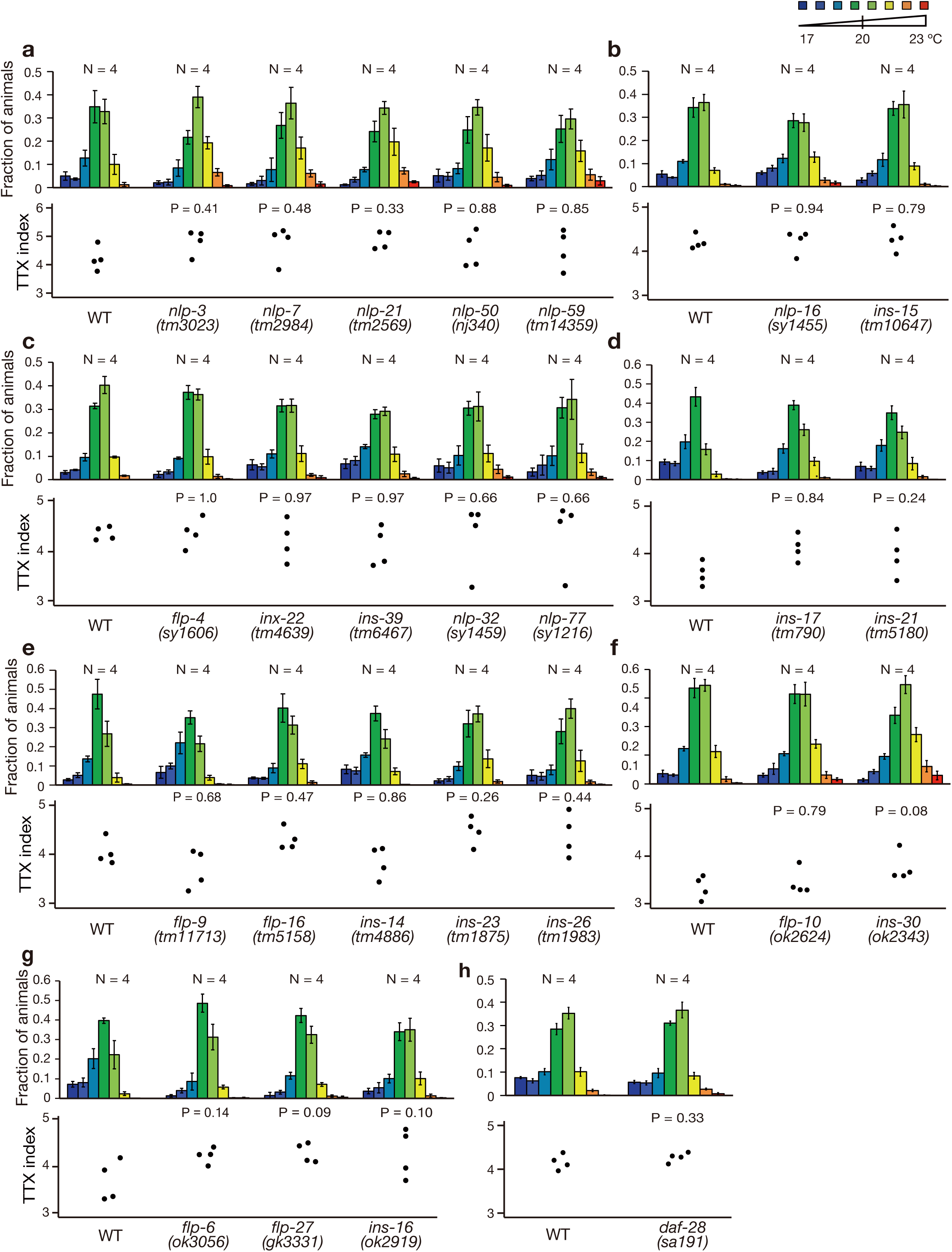
Mutations in the neuropeptide genes expressed in the AFD thermosensory neuron did not affect thermotaxis behavior. (a-h) TTX behaviors of animals in which a neuropeptide gene expressed in the AFD thermosensory neuron is mutated. Distributions of the animals on the thermal gradients of the TTX plate are displayed in the top histograms. The fraction of the population in each temperature section is shown as mean ± SEM. TTX indexes are shown in the bottom dot plots. P-values were determined by Dunnet test in (a), (d), (e) and (g), by Steel test in (b), (c) and (f) or by Student’s T test in (h). Note that *inx-15(tm10647)* animals carry *atm-1(tm5027), xpc-1(tm3886)* and *tm10648,* in their genetic background, and *flp-27(gk3331)* animals *Y17G7B.22(gk1062)* and *gkDf45*.

## References

1. Katz, B. & Miledi, R. Ionic requirements of synaptic transmitter release. Nature 215, 651 (1967).

2. Sabatini, B. L. & Regehr, W. G. Timing of neurotransmission at fast synapses in the mammalian brain. Nature 384, 170–172 (1996).

3. Strathern, L. Getting graded: Teaching principles of chemical synaptic transmission without action potentials. J Undergrad Neurosci Educ 19, R28–R30 (2021).

4. Bennett, M. V. L. & Zukin, R. S. Electrical coupling and neuronal synchronization in the mammalian brain. Neuron 41, 495–511 (2004).

5. Connors, B. W. & Long, M. A. Electrical synapses in the mammalian brain. Annu Rev Neurosci 27, 393–418 (2004).

6. Phelan, P. Innexins: members of an evolutionarily conserved family of gap-junction proteins. Biochim Biophys Acta (BBA) 1711, 225–245 (2005).

7. Söhl, G., Maxeiner, S. & Willecke, K. Expression and functions of neuronal gap junctions. Nat Rev Neurosci 6, 191–200 (2005).

8. Chever, O., Lee, C.-Y. & Rouach, N. Astroglial connexin43 hemichannels tune basal excitatory synaptic transmission. J. Neurosci. 34, 11228–11232 (2014).

9. Cheung, G. et al. Physiological synaptic activity and recognition memory require astroglial glutamine. Nat Commun 13, 753 (2022).

10. Long, M. A., Deans, M. R., Paul, D. L. & Connors, B. W. Rhythmicity without synchrony in the electrically uncoupled inferior olive. J Neurosci 22, 10898–10905 (2002).

11. Leznik, E. & Llinás, R. Role of gap junctions in synchronized neuronal oscillations in the inferior olive. J Neurophysiol 94, 2447–2456 (2005).

12. Dugué, G. P. et al. Electrical coupling mediates tunable low-frequency oscillations and resonance in the cerebellar Golgi cell network. Neuron 61, 126–139 (2009).

13. Yaeger, D. B. & Trussell, L. O. Auditory Golgi cells are interconnected predominantly by electrical synapses. J Neurophysiol 116, 540–551 (2016).

14. Hedgecock, E. M. & Russell, R. L. Normal and mutant thermotaxis in the nematode *Caenorhabditis elegans*. Proc Natl Acad Sci U S A 72, 4061–4065 (1975).

15. Mori, I. & Ohshima, Y. Neural regulation of thermotaxis in *Caenorhabditis elegans*. Nature 376, 344–348 (1995).

16. Luo, L. et al. Bidirectional thermotaxis in *Caenorhabditis elegans* is mediated by distinct sensorimotor strategies driven by the AFD thermosensory neurons. Proc Natl Acad Sci U S A 111, 2776–2781 (2014).

17. Ikeda, M. et al. Context-dependent operation of neural circuits underlies a navigation behavior in *Caenorhabditis elegans*. Proc Natl Acad Sci U S A 117, 6178–6188 (2020).

18. White, J. G., Southgate, E., Thomson, J. N. & Brenner, S. The structure of the nervous system of the nematode *Caenorhabditis elegans*. Philos Trans R Soc Lond B Biol Sci 314, 1–340 (1986).

19. Cook, S. J. et al. Whole-animal connectomes of both *Caenorhabditis elegans* sexes. Nature 571, 63–71 (2019).

20. Kuhara, A. et al. Temperature sensing by an olfactory neuron in a circuit controlling behavior of *C. elegans*. Science 320, 803–807 (2008).

21. Beverly, M., Anbil, S. & Sengupta, P. Degeneracy and neuromodulation among thermosensory neurons contribute to robust thermosensory behaviors in *Caenorhabditis elegans*. J Neurosci 31, 11718–11727 (2011).

22. Kimura, K. D., Miyawaki, A., Matsumoto, K. & Mori, I. The *C. elegans* thermosensory neuron AFD responds to warming. Curr Biol 14, 1291–1295 (2004).

23. Clark, D. A., Biron, D., Sengupta, P. & Samuel, A. D. T. The AFD sensory neurons encode multiple functions underlying thermotactic behavior in *Caenorhabditis elegans*. J Neurosci 26, 7444–7451 (2006).

24. Ramot, D., MacInnis, B. L. & Goodman, M. B. Bidirectional temperature-sensing by a single thermosensory neuron in *C. elegans*. Nat Neurosci 11, 908–915 (2008).

25. Kobayashi, K. et al. Single-cell memory regulates a neural circuit for sensory behavior. Cell Rep 14, 11–21 (2016).

26. Takeishi, A. et al. Receptor-type guanylyl cyclases confer thermosensory responses in *C. elegans*. Neuron 90, 235–244 (2016).

27. Narayan, A., Laurent, G. & Sternberg, P. W. Transfer characteristics of a thermosensory synapse in *Caenorhabditis elegans*. Proc Natl Acad Sci U S A 108, 9667–9672 (2011).

28. Ohnishi, N., Kuhara, A., Nakamura, F., Okochi, Y. & Mori, I. Bidirectional regulation of thermotaxis by glutamate transmissions in *Caenorhabditis elegans*. EMBO J 30, 1376–1388 (2011).

29. Hawk, J. D. et al. Integration of plasticity mechanisms within a single sensory neuron of *C. elegans* actuates a memory. Neuron 97, 356–367.e4 (2018).

30. Nakano, S. et al. Presynaptic MAST kinase controls opposing postsynaptic responses to convey stimulus valence in *Caenorhabditis elegans*. Proc Natl Acad Sci U S A 117, 1638–1647 (2020).

31. Tsukamoto, S. et al. The *Caenorhabditis elegans* INX-4/Innexin is required for the fine-tuning of temperature orientation in thermotaxis behavior. Genes Cells 25, 154–164 (2020).

32. Brenner, S. The genetics of *Caenorhabditis elegans*. Genetics 77, 71–94 (1974).

33. Starich, T. A., Miller, A., Nguyen, R. L., Hall, D. H. & Shaw, J. E. The *Caenorhabditis elegans* innexin INX-3 is localized to gap junctions and is essential for embryonic development. Dev Biol 256, 403–417 (2003).

34. *C. elegans* Deletion Mutant Consortium. large-scale screening for targeted knockouts in the *Caenorhabditis elegans* genome. G3 (Bethesda) 2, 1415–1425 (2012).

35. Chu, J. S.-C. et al. High-throughput capturing and characterization of mutations in essential genes of *Caenorhabditis elegans*. BMC Genomics 15, 361 (2014).

36. Bhattacharya, A., Aghayeva, U., Berghoff, E. G. & Hobert, O. Plasticity of the Electrical Connectome of *C. elegans*. Cell 176, 1174–1189.e16 (2019).

37. Yeh, E., et al. *Caenorhabditis elegans* innexins regulate active zone differentiation. J Neurosci 29, 5207–5217 (2009).

38. Zhang, L., Ward, J. D., Cheng, Z. & Dernburg, A. F. The auxin-inducible degradation (AID) system enables versatile conditional protein depletion in *C. elegans*. Development 142, 4374–4384 (2015).

39. Witvliet, D. et al. Connectomes across development reveal principles of brain maturation. Nature 596, 257–261 (2021).

40. Phelan, P. & Starich, T. A. Innexins get into the gap. Bioessays 23, 388–396 (2001).

41. Oshima, A., Tani, K. & Fujiyoshi, Y. Atomic structure of the innexin-6 gap junction channel determined by cryo-EM. Nat Commun 7, 13681 (2016).

42. Burendei, B. et al. Cryo-EM structures of undocked innexin-6 hemichannels in phospholipids. Sci Adv 6, eaax3157 (2020).

43. Bouhours, M. et al. A co-operative regulation of neuronal excitability by UNC-7 innexin and NCA/NALCN leak channel. Mol Brain 4, 16 (2011).

44. Bai, D., Yue, B. & Aoyama, H. Crucial motifs and residues in the extracellular loops influence the formation and specificity of connexin docking. Biochim Biophys Acta (BBA) 1860, 9–21 (2018).

45. Panchin, Y. et al. A ubiquitous family of putative gap junction molecules. Curr Biol 10, R473–474 (2000).

46. Baranova, A. et al. The mammalian pannexin family is homologous to the invertebrate innexin gap junction proteins. Genomics 83, 706–716 (2004).

47. Beyer, E. C. & Berthoud, V. M. Gap junction gene and protein families: Connexins, innexins, and pannexins. Biochim Biophys Acta (BBA) 1860, 5–8 (2018).

48. Starich, T. A., Xu, J., Skerrett, I. M., Nicholson, B. J. & Shaw, J. E. Interactions between innexins UNC-7 and UNC-9 mediate electrical synapse specificity in the *Caenorhabditis elegans* locomotory nervous system. Neural Dev 4, 16 (2009).

49. Walker, D. S. & Schafer, W. R. Distinct roles for innexin gap junctions and hemichannels in mechanosensation. eLife 9, e50597 (2020).

50. Chelur, D. S. & Chalfie, M. Targeted cell killing by reconstituted caspases. Proc Natl Acad Sci U S A 104, 2283–2288 (2007).

51. Matsuyama, H. J. & Mori, I. Neural coding of thermal preferences in the nematode *Caenorhabditis elegans*. eNeuro 7, ENEURO.0414-19.2020 (2020).

52. Swierczek, N. A., Giles, A. C., Rankin, C. H. & Kerr, R. A. High-throughput behavioral analysis in *C. elegans*. Nat Methods 8, 592–598 (2011).

53. Lee, R. Y., Sawin, E. R., Chalfie, M., Horvitz, H. R. & Avery, L. EAT-4, a homolog of a mammalian sodium-dependent inorganic phosphate cotransporter, is necessary for glutamatergic neurotransmission in *Caenorhabditis elegans*. J Neurosci 19, 159–167 (1999).

54. Speese, S. et al. UNC-31 (CAPS) is required for dense-core vesicle but not synaptic vesicle exocytosis in *Caenorhabditis elegans*. J Neurosci 27, 6150–6162 (2007).

55. Cao, J. et al. Comprehensive single-cell transcriptional profiling of a multicellular organism. Science 357, 661–667 (2017).

56. Taylor, S. R. et al. Molecular topography of an entire nervous system. Cell 184, 4329–4347.e23 (2021).

57. Sandow, S. L. & Hill, C. E. Incidence of myoendothelial gap junctions in the proximal and distal mesenteric arteries of the rat is suggestive of a role in endothelium-derived hyperpolarizing factor-mediated responses. Circ Res 86, 341–346 (2000).

58. Dora, K. A. et al. Myoendothelial gap junctions may provide the pathway for EDHF in mouse mesenteric artery. J Vasc Res 40, 480–490 (2003).

59. Sordillo, A. & Bargmann, C. I. Behavioral control by depolarized and hyperpolarized states of an integrating neuron. eLife 10, e67723 (2021).

60. Chen, G. et al. Connexin-43 induces chemokine release from spinal cord astrocytes to maintain late-phase neuropathic pain in mice. Brain 137, 2193–2209 (2014).

61. Ma, Z. et al. CALHM3 is essential for rapid ion channel-mediated purinergic neurotransmission of GPCR-mediated tastes. Neuron 98, 547–561.e10 (2018).

62. Gray, J. M., Hill, J. J. & Bargmann, C. I. A circuit for navigation in *Caenorhabditis elegans*. Proc Natl Acad Sci U S A 102, 3184–3191 (2005).

63. Kocabas, A., Shen, C.-H., Guo, Z. V. & Ramanathan, S. Controlling interneuron activity in *Caenorhabditis elegans* to evoke chemotactic behaviour. Nature 490, 273–277 (2012).

64. Li, Z., Liu, J., Zheng, M. & Xu, X. Z. S. Encoding of both analog- and digital-like behavioral outputs by one *C. elegans* interneuron. Cell 159, 751–765 (2014).

65. Mello, C. C., Kramer, J. M., Stinchcomb, D. & Ambros, V. Efficient gene transfer in *C. elegans*: extrachromosomal maintenance and integration of transforming sequences. EMBO J 10, 3959–3970 (1991).

66. Dokshin, G. A., Ghanta, K. S., Piscopo, K. M. & Mello, C. C. Robust genome editing with short single-stranded and long, partially single-stranded DNA donors in *Caenorhabditis elegans*. Genetics 210, 781–787 (2018).

67. Ito, H., Inada, H. & Mori, I. Quantitative analysis of thermotaxis in the nematode *Caenorhabditis elegans*. J Neurosci Methods 154, 45–52 (2006).

68. Watanabe, M., Sawada, R., Aramaki, T., Skerrett, I. M. & Kondo, S. The physiological characterization of connexin41.8 and connexin39.4, which are involved in the striped pattern formation of zebrafish. J Biol Chem 291, 1053–1063 (2016).

69. Higurashi, S. et al. Bacterial diet affects the age-dependent decline of associative learning in *Caenorhabditis elegans*. eLife 12, e81418 (2023).

70. Nakano, S. et al. Genetic screens identified dual roles of microtubule-associated serine threonine kinase and CREB within a single thermosensory neuron in the regulation of *Caenorhabditis elegans* thermotaxis behavior. G3 (Bethesda) 12, jkac248 (2022).

